# The use of hybrid data-dependent and -independent acquisition spectral libraries empower dual-proteome profiling

**DOI:** 10.1101/2020.05.24.113340

**Authors:** Patrick Willems, Ursula Fels, An Staes, Kris Gevaert, Petra Van Damme

## Abstract

In the context of bacterial infections, it is imperative that physiological responses can be studied in an integrated manner, meaning a simultaneous analysis of both the host and the pathogen responses. To improve the sensitivity of detection, data-independent acquisition (DIA) based proteomics was found to outperform data-dependent acquisition (DDA) workflows in identifying and quantifying low abundant proteins. Here, by making use of representative bacterial pathogen/host proteome samples, we report an optimized hybrid library generation workflow for data-independent acquisition mass spectrometry relying on the use of data-dependent and *in silico* predicted spectral libraries. When compared to searching DDA experiment-specific libraries only, the use of hybrid libraries significantly improved peptide detection to an extent suggesting that infection relevant host-pathogen conditions could be profiled in sufficient depth without the need of a priori bacterial pathogen enrichment when studying the bacterial proteome.

**GRAPHICAL ABSTRACT:** 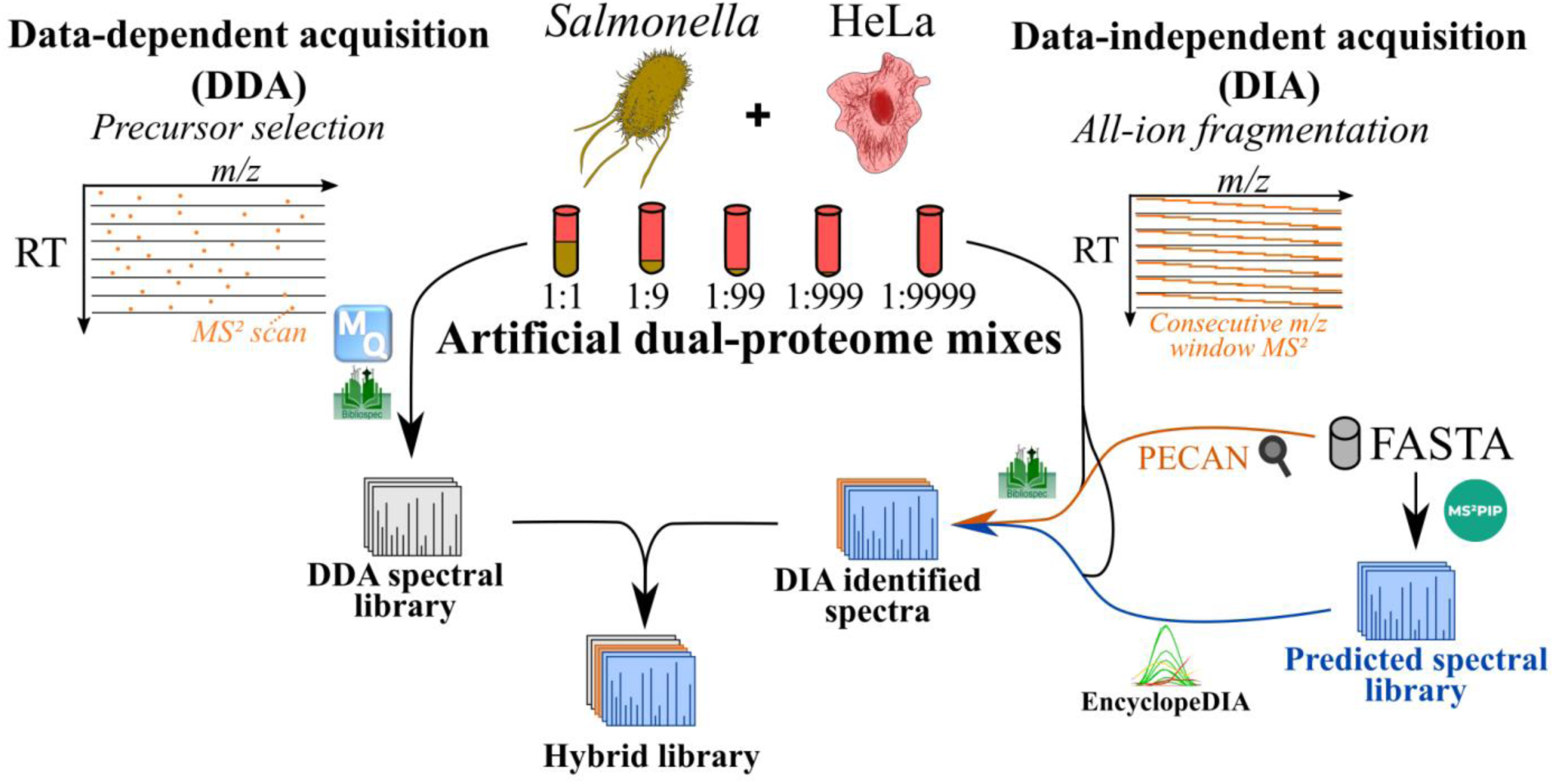

## INTRODUCTION

Amongst others, proteomics aims to identify and quantify changes in protein levels. Bearing in mind that relatively small genomes of microorganism such as that of the bacterium *Salmonella* encode for approximately 4,500 protein-coding genes, bacterial proteomics can be complex, as already a significant number of proteins need to be profiled. In fact, these numbers get even much higher when considering proteoforms, i.e. multiple protein products arising from a single gene [1, 2].

The still preferred method to analyze proteomes is by means of liquid chromatography coupled to tandem mass spectrometry (LC-MS/MS). Specifically, in bottom-up proteomic approaches, proteins are digested to peptides by proteases (e.g. trypsin) which are subsequently analyzed by LC-MS/MS. Mass spectrometers are then mostly operated in so-called data-dependent acquisition (DDA) mode. Here, following an MS1 scan, a number of peptide ions of sufficiently high intensity are selected for further fragmentation to deliver tandem MS (MS2) spectra for sequence identification following database searching [3]. Peptide quantification in DDA is routinely based on the MS1 intensity of the precursor ion, which can be hampered by interference with chemical noise, resulting in a decreased dynamic range due to difficulties when trying to quantify low abundant peptides [4, 5]. On the other hand, in so-called targeted proteomic approaches, e.g. selected reaction monitoring (SRM) and parallel reaction monitoring (PRM), quantifications are based on MS2 scans that are composed of multiple fragments ions, therefore being more robust and more reproducible [6]. However, in this way only a limited number of peptides can be quantified, and thus not permit discovery proteomics. In contrast, mass spectrometers operating in data-independent acquisition (DIA) mode aim to combine the broad identification power of DDA with accurate quantification available by targeted approaches [7]. In DIA mode, all precursors in a predetermined *m*/*z* isolation window are fragmented together irrespective of their abundances. This overcomes the stochastic and thus irreproducible nature of DDA which biases DDA towards the identification of (highly) abundant peptides [4]. Furthermore, in DIA mode, peptide quantification is performed at the MS2 level, resulting in more robust quantification versus the interference-prone MS1 quantification in DDA mode. As the direct relation between precursor and fragment ions is lost, analysis of DIA data requires however very sophisticated algorithms compared to the analysis of DDA data [8].

While spectrum-centric DDA search algorithms are not suited for the analysis of the complex MS2 spectra generated by DIA, annotated MS2 spectra and retention times (RTs) from DDA searches are widely used for constructing spectral libraries to query DIA data [9–11]. The sensitivity of these approaches is thus inherently limited to the stochastic limitations of DDA. Spectral libraries can be created in several ways, leading to more or less extensive libraries, with the library size influencing the accuracy and specificity of identifications and quantifications [12]. Besides the use of DDA-based spectral libraries, an alternative way of creating spectral libraries is by using algorithms that accurately predict MS2 fragmentation spectra such as MS^2^PIP [13] and Prosit [14]. MS^2^PIP (MS2 peak intensity prediction) is a data-driven tool [13] that is trained on different types of public DDA data to predict MS2 peak intensities [15]. Prosit [14] on the other hand is trained on MS2 spectra of synthetic peptides and mass spectrometry data generated in the context of the ProteomeTools project, which has the overall aim to provide a high-quality reference MS2 data of synthetic peptides [16]. In this way, Prosit predicts fragment ion intensities and RTs for a given peptide-charge pair and can generate spectral libraries ready for DIA searches and, recently, a graphical user interface was built in the EncyclopeDIA library search engine [17]. Next to predicted spectral libraries, so-called library-free approaches overcome the stochastic limitation of DDA by querying DIA data for the best supporting evidence of peptide detection. The peptide-centric algorithm <u>P</u>eptide <u>C</u>entric <u>A</u>nalysis or PECAN [18] incorporates a sequence-based RT predictor to improve the sensitivity of detection by filtering DIA data based on expected RTs. In a DIA-only workflow, PECAN uses as input DIA data, a queried peptide list with a background proteome and provides as output auxiliary scores of experimental and expected RTs, which are further processed by Percolator [19] to report confident peptide and protein identifications. Another library-free approach include algorithms such as DIA-Umpire that perform deconvolution of DIA data to DDA-like pseudo spectra, which can then be identified with DDA-based database searching approaches [20].

In bacterial infection biology, a comprehensive mapping of the proteome profiles of both host and pathogen is needed [21]. Such dual proteome profiles are currently lacking, likely because of the scarce amount of protein material originating from the bacterial pathogen in most commonly used infection models. Since most intracellular bacteria replicate inside the host, bacteria enumeration varies during the time course of an infection. More specifically, in *in vitro* infection models using *Salmonella* infected epithelial HeLa cells, bacterial enumeration estimates <10 bacteria per HeLa cell at early times and up to 100 bacteria at late times in infection[22]. Considering these numbers, bacteria to host protein content varies from ~1:1000 to 1:100 during the infection process. Despite the improved sensitivity and speed of contemporary mass spectrometers, the current technologies still do not allow to study host and pathogen proteomes simultaneously with sufficient depth, requiring prior enrichment pf bacteria which can typically be attained by selective host lysis and differential centrifugation [23, 24]. Moreover, as from the host point of view, bystander cells can first be removed in order to only profile the infected host cells’ proteomes. This is commonly done by using fluorescent reporter strains and fluorescenc activated cell sorting (FACS) [25, 26]. Of note, the elimination of bystander contributions, concomitantly enriches for bacterial content as a typically only a fraction of cells get infected [27].

In the present work, we aimed to establish a DIA-MS workflow to improve the overall sensitivity of protein identification and quantification of complex *Salmonella*-host mixtures containing only a fairly low amount of peptide material derived from *Salmonella* without bacterial pre-enrichment. We compared the performance of MS^2^PIP-predicted and DDA-based spectral libraries, and, to improve on DDA sensitivity, we extended the spectral library [28] by including MS2 spectra obtained from pre-fractionated LC-MS/MS DDA data of *Salmonella* grown under different infection-relevant conditions. Lastly, by integrating (predicted) DDA libraries and library-independent approaches, a hybrid DDA-based spectral library was created [29], which could achieve an up to 2-to 3-fold increase in consistently quantified human or spiked in *Salmonella* proteins and peptides.

## EXPERIMENTAL PROCEDURES

### HeLa cell culture

HeLa cells (epithelial cervix adenocarcinoma, American Type Culture Collection, Manassas, VA, USA; ATCC® CCL-2™) were cultured in GlutaMAX containing Dulbecco’s Modified Eagle Medium (DMEM) (Gibco, cat n° 31966047) supplemented with 10% fetal bovine serum (Gibco, cat n° 10270-106) and 50 units/mL penicillin and 50 µg/mL streptomycin (Gibco; cat n° 5070-063). Cells were cultured at 37°C in a humidified atmosphere with 5% CO_2_ and passaged at an 1:8 ratio every 4 days.

### Bacterial strain and *Salmonella* cultivation conditions

The *Salmonella enterica* serovar Typhimurium wild-type strain SL1344 [30] (Genotype: hisG46, Phenotype: His(-); biotype 26i), herein referred to as *Salmonella*, was obtained from the *Salmonella* Genetic Stock Center (SGSC, Calgary, Canada; cat n° 438). Bacterial growth was performed in liquid Lennox (L) growth medium (10 g/L Bacto tryptone, 5 g/L Bacto yeast extract, 5 g/L NaCl), Luria Beltrami (LB)-Miller broth (10 g/L Bacto tryptone, 5 g/L Bacto yeast extract, 10 g/L NaCl) or variants of phosphate carbon nitrogen (PCN) medium [31] (lnSPI2; pH 5.8, 0.4 mM Pi), SPI2-inducing PCN (pH 5.8, 0.4 mM inorganic phosphate) containing low levels (10 μM) of magnesium sulfate (PCN medium was stored at 4 °C and brought at room temperature for cultivation). Viewing the auxotrophic nature of the SL1344 strain used, all PCN media were supplemented with histidine to a final concentration of 5 mM. For bacterial cultivation, single colonies were picked from LB plates, inoculated in 8 mL liquid Lennox (L) growth medium (L-broth) and grown overnight at 37 °C with agitation (180 rpm). Subsequently, the overnight cultures with an optical density measured at 600 nm (OD_600_) of ~4.8 were diluted 1:200 (~OD_600_ 0.02) in T175 flasks in 50 mL L-medium without antibiotics and grown under ten different (infection-relevant) growth conditions as reported in [32]. More specifically, bacteria were grown to early exponential growth phase (EEP; OD_600_ 0.1), mid exponential growth phase (MEP; OD_600_ 0.3), late exponential growth phase (LEP; OD_600_ 1.0), early stationary phase (ESP; OD_600_ 2.0) and late stationary phase (LSP; OD_600_ 2.0 + 6 h of extra growth). Besides, environmental shocks in LB were performed on MEP-grown bacteria by the addition of NaCl to a final concentration of 0.3 M and continued growth for 10 min or, in case of anaerobic shock, growth for an additional 30 min without agitation in a filled and tightly screwed 50 mL Falcon tube. For growth in variants of PCN minimal medium [31], overnight grown LB cultures were washed twice in PCN medium before resuspension at O.D600 0.02. Cells were grown in SPI2-inducing PCN or low magnesium SPI2-inducing PCN. The nitric oxide shock conducted in PCN (InSPI2) was performed at OD_600_ 0.3 by addition of the nitric oxide donor spermine NONOate to a final concentration of 250 μM for 20 min (nitric oxide shock (InSPI2)) [33]. Bacterial cells were collected by centrifugation (2,600 × *g*, 10 min) at 4 °C and the supernatant discarded. Samples were flash frozen in liquid nitrogen and stored at −80 °C until further processing.

### Proteome extractions and sample preparation

Cell pellets were resuspended in guanidinium chloride (Gu.HCl) containing lysis buffer (4 M Gu.HCl, 50 mM ammonium bicarbonate (pH 7.9)) at 5 x 10^9 *Salmonella* and 1 x 10^7 Hela cells per 500 µL lysis buffer and mechanically lysed by three rounds of freeze-thaw cycles in liquid nitrogen. The lysates were sonicated (Branson probe sonifier output 4, 50% duty cycle, 2×30 s, 1 s pulses) followed by centrifugation (16,100 x *g*, 10 min) at 4 °C, to remove cellular debris. The protein concentration of the supernatant was determined by Bradford measurement according to the manufacturer’s instructions (Bio-Rad, cat n° 5000006).

Samples were made from trypsin digested total *Salmonella* and/or HeLa protein lysate(s) (mixtures). More specifically, for the infection-relevant complex *Salmonella* sample, an equimolar mix of *Salmonella* protein samples originating from *Salmonella* grown in the 10 infection-relevant conditions was made. Alternatively, in case of complex *Salmonella*-host mixtures, *Salmonella* protein lysate (S) dilution series in protein lysates of human HeLa cells (H) (hereafter referred to as artificial mixtures), were made by mixing proteome samples prior to digestion in a 1:9 ratio, and making dilutions series thereof to obtain complex S:H proteome mixtures with corresponding *Salmonella*/HeLa protein ratios of 1:99, 1:999 and 1:9999. In addition, a sample containing equal amounts of *Salmonella* and HeLa proteins (1:1 ratio) was prepared. All spiked in samples were prepared in triplicate. For all protein mixtures, an aliquot equivalent to 400 µg of total protein was transferred to a 1.5 ml Eppendorf tube, twice diluted with HPLC-grade water and precipitated overnight with 4 volumes of −20 °C acetone. The precipitated protein material was recovered by centrifugation (3,500 x *g*, 15 min) at 4 °C, pellets were washed twice with −20°C 80% acetone, and air dried upside down at room temperature until no residual acetone odor remained. Pellets were resuspended in 200 µL TFE (2,2,2-trifluoroethanol) digestion buffer (10% TFE, 100 mM ammonium bicarbonate, pH 7.9) with sonication at 4°C (Branson probe output 20; 1 s pulses) until a homogenous suspension was reached. All samples were digested overnight at 37 °C using a Trypsin/Lys-C Mix (mass spec grade, Promega, Madison, WI) (enzyme/substrate of 1:100 w/w) while mixing (550 rpm). Samples were acidified with TFA to a final concentration of 0.5%, cleared from insoluble particulates by centrifugation (16,100 x *g*, 15 min) at 4 °C and the supernatant transferred to new Eppendorf tubes. Methionine oxidation was performed by the addition of hydrogen peroxide to a final concentration of 0.5% for 30 min at 30 °C. Solid phase extraction of peptides was performed using C18 reversed phase sorbent containing 100 µL pipette tips according to the manufacturer’s instructions (Agilent, Santa Clara, CA, USA, cat n° A57003100K). The pipette tip was conditioned by aspirating the maximum pipette tip volume of water:acetonitrile, 50:50 (v/v) and the solvent was discarded. After equilibration of the tip by washing 3 times with the maximum pipette tip volume in 0.1% TFA in water, 100 µl of the acidified peptide mixtures (~200 µg) was dispensed and aspirated for 10 cycles for maximum binding efficiency. The tip was washed 3 times with the maximum pipette tip volume of 0.1% TFA in water:acetonitrile, 98:2 (v/v) and the bound peptides were eluted in LC-MS/MS vials with the maximum pipette tip volume of 0.1% TFA in water:acetonitrile, 30:70 (v/v). The samples were vacuum-dried in a SpeedVac concentrator and re-dissolved in 100 µL (infection-relevant *Salmonella* peptide mixture) for subsequent RP-HPLC fractionation (see below) or, for LC-MS/MS analysis, in 50 µL (artificial mixtures) or 20 µL (fractionated RP-HPLC samples obtained from the infection-relevant *Salmonella* peptide mixture) of 2 mM tris(2-carboxyethyl)phosphine (TCEP) in 2% acetonitrile spiked with an indexed Retention Time (iRT) peptide mix (Biognosys) according to the manufacturer’s instructions [34] for retention time (RT) prediction (see below). Samples were stored at −20 °C until further analysis.

### RP-HPLC fractionation of complex *Salmonella* peptide mixture

The infection-relevant, complex *Salmonella* peptide mixture (100 µl) was acidified by the addition of 5 µl of glacial acetic acid and the peptide mixture (corresponding to an equivalent of 400 µg of digested protein) fractionated at pH 5 using a High Pressure Liquid Chromatography (HPLC) Agilent series 1100 instrument. The sample was trapped for 16 min on a reversed-phase trapping column (35 mm x 300 µm I.D., 5 µm beads C18 material (Dr. Maisch, Germany), fritted and packed in-house). Next, the sample was separated on an analytical column (150 mm x 250 µm I.D., 3 µm beads C18 material (Dr. Maisch, Germany), fritted and packed in-house) using a 100 min gradient from solvent A (10 mM ammonium acetate, pH 5.5) to solvent B (10 mM ammonium acetate, 70% ACN, pH 5.5) at a constant flow rate of 3 µl/min. The constant flow rate is achieved with Agilent’s 1100 series capillary pump in micro flow mode with the flow controller at 20 µl/min. After the gradient, the column was run at solvent B for 5 min, switched to solvent A and re-equilibrated for 20 min. One-minute fractions were collected in MS vials over a time interval of 65 min and automatically pooled, this being the restarting of the fraction collection cycle every 10 min resulting in a total of 10 (pooled) fractions. Peptide were detected by absorbance at 214 nm and 280 nm. The fractions were then vacuum-dried in a SpeedVac concentrator and re-dissolved in 20 µL of 2 mM TCEP in 2% acetonitrile spiked with the iRT peptide mix as described above. Samples, referred to as *Salmonella* pre-fractionated samples, were stored at −20°C until further analysis.

### LC-MS/MS data acquisition

From each artificial mixture and *Salmonella* pre-fractionated sample, 10 µL and 2 µL was injected onto the column, corresponding to 2 µg and 4 µg peptide material, respectively for LC-MS/MS analysis on an Ultimate 3000 RSLC nano system in-line connected to a Q Exactive HF BioPharma mass spectrometer (Thermo). Trapping was performed at 10 μl/min for 4 min in loading solvent A (0.1% TFA in water/ACN (98:2, v/v) on a 20 mm trapping column (made in-house, 100 μm internal diameter (I.D.), 5 μm beads, C18 Reprosil-HD, Dr. Maisch, Germany). After flushing from the trapping column, the peptides were loaded and separated on an analytical 200 cm µPAC™ column with C18-endcapped functionality (PharmaFluidics, Belgium) kept at a constant temperature of 50 °C. Peptides were eluted using a non-linear gradient reaching 9% MS solvent B (0.1% formic acid (FA) in water/acetonitrile (2:8, v/v)) in 15 min, 33% MS solvent B in 105 min, 55% MS solvent B in 125 min and 99% MS solvent B in 135 min at a constant flow rate of 300 nL/min, followed by a 5-min wash with 99% MS solvent B and re-equilibration with MS solvent A (0.1% FA in water). For both analyses, a pneu-Nimbus dual column ionization source was used (Phoenix S&T), at a spray voltage of 2.6 kV and a capillary temperature of 275 °C. For the first analysis (DDA mode), the mass spectrometer automatically switched between MS and MS2acquisition for the 16 most abundant ion peaks per MS spectrum. Full-scan MS spectra (375-1500 *m*/*z*) were acquired at a precursor resolution of 60,000 at 200 *m*/*z* in the Orbitrap analyser after accumulation to a target value of 3,000,000. The 16 most intense ions above a threshold value of 13,000 were isolated for higher-energy collisional dissociation (HCD) fragmentation at a normalized collision energy of 28% after filling the trap at a target value of 100,000 for maximum 80 ms injection time using a dynamic exclusion of 12 s. MS2 spectra (200-2,000 *m*/*z*) were acquired at a resolution of 15,000 at 200 *m*/*z* in the Orbitrap analyser.

Another 10 µL aliquot from each artificial mixture was analysed using the same mass spectrometer in DIA mode. Nano LC conditions and gradients were the same as used for DDA. Full-scan MS spectra ranging from 375-1,500 *m*/*z* with a target value of 5E6 were followed by 30 quadrupole isolations with a precursor isolation width of 10 *m*/*z* for HCD fragmentation at a normalized collision energy of 30% after filling the trap at a target value of 3E6 for maximum injection time of 45 ms. MS2 spectra were acquired at a resolution of 15,000 at 200 *m*/*z* in the Orbitrap analyser without multiplexing. The isolation intervals ranged from 400 – 900 *m*/*z* with an overlap of 5 *m*/z.

### Processing of data-dependent acquisition (DDA) data

Raw data files corresponding to 10 fractions of the complex, infection-relevant *Salmonella* peptide mixture and 5 artificial dual-proteome mixtures (1:1, 1:9, 1:99, 1:999 and 1:9999 *Salmonella*:HeLa samples, in triplicates) were searched in parallel by MaxQuant [35] (version 1.6.10.43). Protein databases for searching the obtained spectra were either the UniProt knowledgebase (UniProtKB) proteomes for *Salmonella* pre-fractionated samples (proteome UP000008962, 4,657 proteins) or the *Salmonella* proteome database concatenated to the human UniProtKB database for artificial mixtures (proteomes UP000008962 [4,657 proteins] and UP000005640 [74,449 proteins]). In addition, MaxQuant built-in contaminant proteins and the 11 iRT peptide sequences (Biognosys-11) were included in the search [34]. Methionine oxidation to methionine-sulfoxide was set as a fixed modification, and in case of artificial mixtures, protein N-terminal acetylation was set as variable modification. To augment peptide quantification, we performed matching-between-runs with a match time window of 1.2 min and an alignment time window of 20 min, and performed label-free quantitation with the LFQ algorithm using default settings in MaxQuant. We used the enzymatic rule of trypsin/P with a maximum of two missed cleavages. Peptide-to-spectrum match level was set at 1% FDR. Protein FDR – calculated by employing a reverse database strategy – was set at 1%. For protein quantification in the proteinGroups.txt file, only unique peptides were considered and all modifications were allowed. For other search parameters not specified here, default MaxQuant settings were used.

### Processing of data-independent acquisition (DIA) data

#### DDA-based spectral library construction

The ‘msms.txt’ files outputted by MaxQuant were used as input for the creation of redundant BLIB spectral libraries (artificial mixtures and *Salmonella* pre-fractionated samples) using BiblioSpec (version 2.1) [36]. Redundant spectra were subsequently filtered using the ‘BlibFilter’ function, requiring entries to have at least 5 peaks (‘-n 5’). Peptide sequences uniquely identified in the *Salmonella* pre-fractionated samples were appended to the spectral library of the artificial mixtures to extend the library and augment the detection capacity of *Salmonella* peptides. We transformed the peptide RTs present in the BLIB library to iRTs using the spiked-in iRT peptides (Biognosys-11). To this end, empirical RTs of the top-scoring iRT peptide identifications (lowest posterior error probability, ‘msms.txt’) in the DDA samples were used to fit a linear trendline and scale the RTs. For the artificial mixtures and *Salmonella* pre-fractionation analyses, the corresponding trendlines were iRT = 1.220 RT – 74.566 and iRT = 1.189 RT – 75.039, respectively. The updated BLIB files were then converted to DLIB format by EncyclopeDIA, using the combined human-*Salmonella* UniProtKB proteome FASTA as background [37].

#### Library-free searching of proteome FASTA by PECAN/Walnut

DIA raw data files were converted to mzML by MSConvert using vendor peakPicking and enabling the ‘SIM as spectra’ option. Pre-processed DIA samples were searched against a compilation of the *Salmonella* UniProtKB proteome (UP000008962, 4,657 proteins) and human Swiss-Prot proteome (UP000005640, 20,367 proteins) using the EncylopeDIA built-in PECAN algorithm [18, 37]. We opted to solely search the human Swiss-Prot protein database, which resulted in a ~3-fold reduction of the protein database search space (25,024 versus 74,449 proteins when combining *Salmonella* [UP000008962] and human [UP000005640] UniProtKB references proteomes), in order to minimize the theoretical peptide search space. This was desired to limit the size of a predicted spectral library for all possible tryptic peptides (see below) and overall runtime and memory usage. Since 36,494 out of 36,668 (i.e. 99.53%) of all human peptides identified by MaxQuant matched a Swiss-Prot protein entry, no drastic loss in identifications is anticipated. Default settings were used, except for methionine oxidation (to methionine-sulfoxide) being set as fixed modification, and considering a maximum length of 25 amino acids and HCD as fragmentation type.

#### Construction of an MS^2^PIP-based spectral library

MS2 spectra were predicted by MS^2^PIP (version 20190312) [13] for tryptic peptides derived from an *in silico* digest of the *Salmonella* UniProtKB proteome and human Swiss-Prot proteome (trypsin/P, peptide length 7-25 AA, mass 500-5,000 Da, one missed cleavage, N-terminal initiator methionine removal considered) in case of 2+ and or 3+ peptide precursor fit within 400 to 900 *m/z* (scanned range DIA). This yields a total of 1,586,777 predicted MS2 spectra for 1,151,386 peptides solely matching human proteins, 197,782 spectra for 144,156 peptides matching *Salmonella* proteins, and 117 spectra for 110 peptides matching both species. We set methionine oxidation (to methionine-sulfoxide) as a fixed modification for MS2 prediction by MS^2^PIP. Predicted spectra were supplemented with DeepLC predicted RTs using a model trained on RTs of 35,206 non-redundant peptides identified in DIA PECAN searches (peptide Q-value < 0.01) (as described in the above section).

#### Hybrid library construction

The chromatogram libraries (ELIB) generated by EncyclopeDIA after PECAN and MS^2^PIP-library searching were combined into a single redundant library. After conversion to BLIB format and adding the ‘redundant’ tag in library info (sqlite3), the ‘BlibFilter’ function was used to create a non-redundant DIA spectral library as described above. This DIA spectral library was further extended with non-redundant spectra from the extended DDA spectral library (see above). The resulting hybrid DDA/DIA spectral library (i.e. joining DDA- and DIA-based identifications) was then converted to DLIB format as described above. The data analysis workflow of hybrid DDA-DIA spectral library construction is provided in Figure 1.

**Figure 1.**
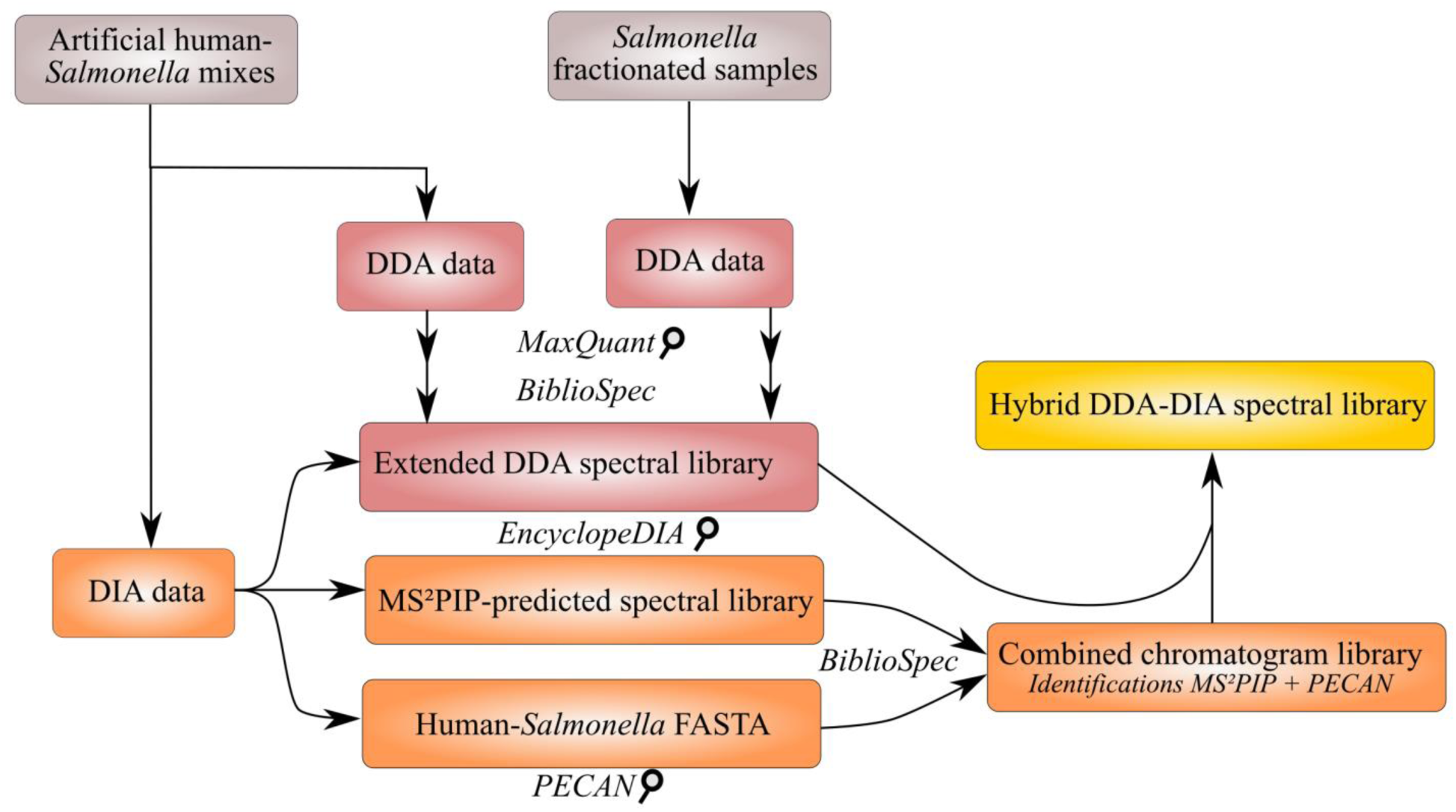
Proteome data analysis workflow of hybrid DDA-DIA spectral library construction. DDA data from artificial dual-proteome mixtures and complex *Salmonella* pre-fractionated samples were searched using MaxQuant [35], and an extended DDA spectral library of both datasets was constructed using BiblioSpec [36]. In parallel, entire proteomes of human and *Salmonella* were searched either by searching an MS^2^PIP-predicted spectral library with EncyclopeDIA [37] or searching the combined FASTA using PECAN [18]. A combined chromatogram library (ELIB) was constructed from the identification results of whole proteome DIA-only searches. Lastly, this combined chromatogram library was appended with DDA spectra which were assigned to peptides not yet present in the chromatogram library to generate a hybrid DDA-DIA spectral library.

#### EncyclopeDIA spectral library searching and peptide quantification

The resulting mzML files were searched against the DDA, MS^2^PIP, or hybrid spectral DLIB libraries using EncylopeDIA software (version 0.90) [37] with default settings. Sample-specific Percolator output files and chromatogram libraries were stored. Per setup, a combined chromatogram library was created consisting of the three replicates. This performs a Percolator re-running of the combined results and provides peptide and protein quantifications at a 1% peptide and protein Q-value, respectively. For quantification, the number of minimum required and quantifiable ions were set at 5 with aligning between samples enabled.

## RESULTS

### DDA-based spectral library searching of DIA data improves detection of low abundant peptides

To mimic the proteome complexity of bacterial host cell infections, we generated five artificial *Salmonella*-human proteome mixtures (1:1, 1:9, 1:99, 1:999 and 1:9999) in triplicate, referred to as artificial dual-proteome mixtures. Following trypsin digestion, each sample was analysed by LC-MS/MS in both DDA and DIA mode. First, DDA data were analysed by MaxQuant against a composite database containing *Salmonella* as well as human UniProtKB protein entries (see Experimental Procedures). As anticipated, with decreasing *Salmonella* protein content, the number of identified *Salmonella* peptides decreases, whereas the number of identified human peptides increases (Figure 2A, orange bars, top to bottom). Indeed, whereas approximately 5,008 non-redundant *Salmonella* peptides are consistently (i.e. in all three replicates) identified in the 1:1 dilution, this number decreases to 1,226 (24.5%) and only 257 (5.1%) in the 1:9 and 1:99 dilutions, respectively.

**Figure 2.**
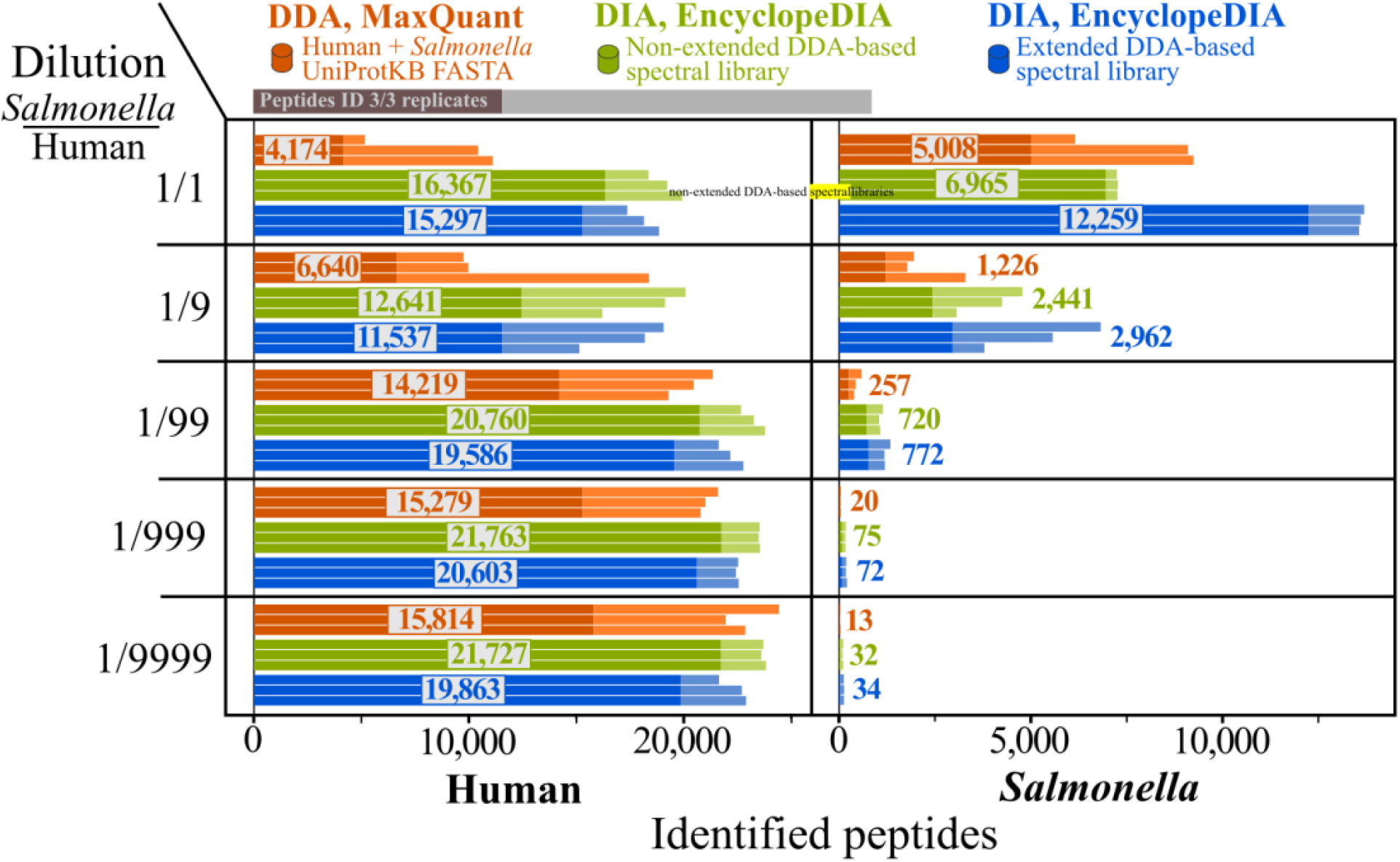
Peptide identifications in artificial dual proteome mixtures acquired in DDA and DIA modes. The number of identified non-redundant human (left panel) or *Salmonella* (right panel) peptide sequences (x-axis, peptide Q-value ≤ 0.01) are shown per replicate across artificial mixtures (y-axis, 1:1 to 1:9999). Bars showing triplicate samples are used to display data acquired in DDA mode and searched with MaxQuant (orange), data acquired in DIA mode and searched with EncyclopeDIA against the non-extended (green) and extended (blue) DDA-based spectral library. The dark-coloured portion of the bars and corresponding numbers within indicate the number of peptide sequences consistently identified in all three replicate samples.

In a next phase, MaxQuant results were used to create a DDA-based spectral library (see Experimental Procedures), encompassing a total of 57,056 MS2 spectra corresponding to 44,932 non-redundant peptide sequences of which 34,475 (76.7%) uniquely matched to human proteins and 10,422 (23.2%) to *Salmonella* proteins. However, when analysing related shotgun samples exclusively consisting of *Salmonella* peptides on the same MS instrument, we previously routinely identified between 15,000 to 25,000 *Salmonella* peptides with MaxQuant [38], suggesting that mixing of human and *Salmonella* proteomes limits *Salmonella* peptide identification and thus DDA-based spectral library construction. To extend the number of *Salmonella* peptides in this library, we performed an offline RP-HPLC pre-fractionation of a digest of a complex proteome mixture obtained from mixing equal proteome amounts from *Salmonella* grown *in vitro* under 10 different (infection relevant) conditions as reported in [32] (MaxQuant results, see Dataset PXD017904). This way, an additional 12,924 non-redundant *Salmonella* peptide sequences (27,575 spectra) were appended to the DDA-based spectral library. These two DDA libraries are herein referred to as non-extended and extended DDA-based spectral libraries, respectively. DIA data from artificial mixtures were searched against both spectral libraries using EncyclopeDIA [37] (Figure 2, green and blue bars, respectively). In case of human peptides, DIA analysis shows a notable increase (up to 3.7-fold) of identifications in the 1:1 and 1:9 dilutions compared to DDA analysis, whereas presenting similar identification numbers (~16,000-18,000 human peptides) when decreasing *Salmonella* peptide input further. Overall, this suggests that with increasing abundance, *Salmonella* peptide ions obstruct selection and fragmentation of human peptide ions in DDA mode, while clearly much less interference is observed when samples are analysed in DIA modus [39]. Further, when looking at artificial mixtures across all dilutions, DIA identifies most peptides consistently (i.e. in all three replicates). Notably, extending the DDA spectral library with additional *Salmonella* peptides (Figure 2, left panel, blue versus green bars) results in a slight decrease of human peptide identifications. Logically, the opposite trends holds true when inspecting *Salmonella* peptide identifications (Figure 2, right panel). For instance, in all dilutions, the DIA data queried with the extended DDA-based spectral library more than doubles the number of consistently identified *Salmonella* peptides compared to DDA identifications. Taken together, offline fractionation methods can provide a useful asset to improve DIA identification rates, especially in case of complex proteome mixes with components of low abundance (e.g. dual proteome mixes such as bacterial pathogen infection of human hosts). Most likely, the multi-species peptide mixture here is impossible to grasp in the given LC-MS/MS setting – illustrated by merely ~ 10,000 identified *Salmonella* peptides, whereas on average more than double *Salmonella* peptides are identified in pure proteome samples.

### DIA-only workflows and predicted spectral libraries can join forces with DDA spectral libraries

While our DIA spectral library searches clearly outperformed MaxQuant DDA analysis in terms of peptide identification, an important limitation is that only those peptides originally identified in DDA analysis can be detected. To further tap into the peptide discovery potential of DIA data, we implemented two workflows independent of DDA data. First, we used the DIA library-free PECAN software to search the human and *Salmonella* proteome. When using PECAN, the number of peptide identifications are in line with or just below DDA data-based identifications but lower when our DIA data was queried with the extended DDA-based spectral library, in line with previous reports [37]. Second, we generated a spectral library of an *in silico* tryptic proteome digest of the human and *Salmonella* proteomes using MS^2^PIP [13]. In total, 1,784,677 MS2 spectra were predicted for 2+/3+ peptide precursors (395 to 905 *m/z*) for 1,295,652 peptides with enzymatic settings similar to PECAN search settings (Trypsin/P, 7 to 25 amino acid long peptides with a maximum of one missed cleavage). DIA-based peptide RTs obtained from the PECAN search results were used to train and predict RTs for all peptides using DeepLC [40]. We then used the MS^2^PIP-predicted spectral library to query our artificial mixtures with EncyclopeDIA (Figure 3, left panel, grey bars). With the exception of the 1:1 samples, the searches identified a similar number of human peptides per replicate, though a lower number of consistently identified peptides was observed among replicates. This relatively lower consistency is likely due to the higher amount of peptide sequences, ~35-fold, in the MS^2^PIP-predicted *in silico* spectral library, when compared to the extended DDA-based spectral library. The use of large database sizes already showed to increase variation in peptide and protein quantification [12]. Importantly, and as illustrated in the respective Venn diagrams (insets Figure 3, left panel), thousands of peptides not present either in the DDA results nor DDA-based spectral libraries were identified using the MS^2^PIP-predicted *in silico* spectral library, demonstrating the potential of this approach to identify peptides in DIA not found in DDA. Turning our attention to *Salmonella* identified peptides, the MS^2^PIP-predicted *in silico* library shows relatively lower peptide identifications rates (~60-80%) than the extended DDA-based spectral library. Nonetheless, a significant agreement among results is observed, as for instance in the 1:1 dilutions, 7,727 out of 8,520 peptide sequences (90.7%) identified in the MS^2^PIP-predicted *in silico* spectral library search were also identified when querying the artificial mixtures with the DDA-based spectral library. In a next phase, we inspected the properties of peptides identified by searching the MS^2^PIP-predicted spectral library that were not found in the DDA searches (and thus not part of the DDA spectral library). For instance, 1,594 peptides were discovered using the MS^2^PIP-predicted spectral library in the 1:1 sample, of which 835 human and 759 *Salmonella* peptides (Supplementary Figure 2). In less diluted human samples 1:999 and 1:9999, more than 4,000 peptides are uniquely found using the MS^2^PIP-predicted spectral library. When comparing the peptide quantifications of peptides included in the DDA-based spectral library and those discovered by the MS^2^PIP-predicted spectral library searches, the novel MS^2^PIP-identified peptide distributions are slightly lower, and yet more outspoken in the 1:999 and 1:9999 samples (Supplementary Figure 2A). Hence, suggesting that the spectral library enables the detection of relatively lower abundant peptide species missed in DDA. Next, we checked whether the novel MS^2^PIP-based peptides matched to protein entriess that were matched by other peptides in the DDA searches. In all dilutions, approximately 85% of MS^2^PIP-based peptides matched a protein identified in the MaxQuant searches of the DDA data (Supplementary Figure 2B). As such, the majority of MS^2^PIP-based peptides increase the protein sequence coverage for proteins present in the DDA spectral library in turn increasing identification and quantification confidence.

**Figure 3.**
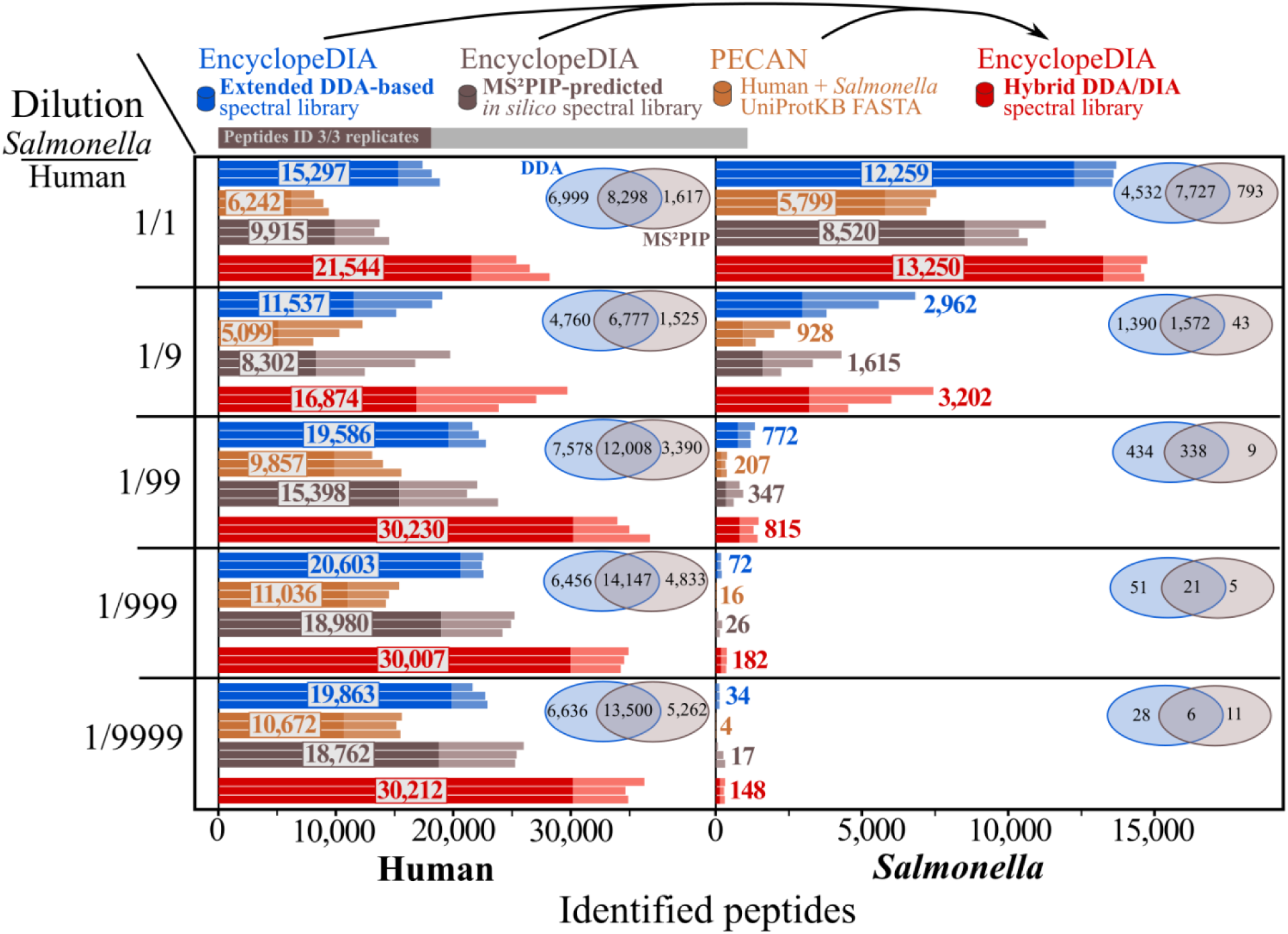
DIA based peptide identifications using different search strategies. The number of identified non-redundant human (left panel) or *Salmonella* (right panel) peptide sequences (x-axis, peptide Q-value ≤ 0.01) are shown per replicate across artificial mixtures (y-axis, 1:1 to 1:9999). Bars showing triplicate samples are used to display data acquired in DIA mode and searched with the extended DDA-based spectral library (blue), the MS^2^PIP-predicted *in silico* spectral library (grey), EncyclopeDIA built-in PECAN algorithm (Walnut) using as input the human-*Salmonella* FASTA (brown), and the hybrid DIA/DDA spectral library encompassing all DIA search results merged with DDA search results (red). The dark-coloured portion of the bars and corresponding numbers within indicate the number of peptide sequences consistently identified in all three samples. Venn diagrams indicate the overlap of consistently identified peptide sequences between the MS^2^PIP-predicted *in silico* spectral library and DDA-based spectral library.

In a next phase, we generated a hybrid spectral library to attest its potential to further increase proteome coverage (Experimental procedures, Figure 1). Hybrid spectral libraries are essentially merged libraries comprising results of different DDA or DIA analysis workflows, e.g. combining predicted spectral libraries with experimental libraries [41], or DDA-based and DIA-only spectral libraries [29]. Here, the MS^2^PIP-predicted spectral library search, PECAN analysis and obtained MaxQuant DDA results were merged to generate a single hybrid library. More specifically, 21,076 DDA-only identified peptide sequences (27,788 spectra) were joined with 55,537 peptide sequences identified in DIA-searches (59,745 spectra), of which 18,736 peptide sequences (33.7%) were solely identified upon DIA and not DDA analysis of the artificial mixtures. The hybrid library resulted in a higher (consistent) number of peptide identifications for both species across all dilutions (Figure 3, red bars). This positive effect was most outspoken for human peptide identifications, where in the 1:99 to 1:9999 samples, ~30,000 peptides were consistently identified in all three replicates, whereas only about 20,000 human peptides were identified when solely considering DDA results.

### DIA improves quantification of low abundant proteins

Besides the improved peptide identification rates, we evaluated and compared protein quantifications across dilutions between both acquisition modes. Aiming at assessing the performance per dilution series independently, i.e. as a proxy for biological infection conditions where the bacterial proteome content is limiting and thus 1:1 and 1:9 dilution conditions are thus typically not representative, we ran the three replicates of each dilution in MaxQuant enabling matching-between-runs and used the alignment-between-runs function in EncyclopeDIA. Importantly, we only considered protein quantifications per dilution series if quantified in at least two out of three replicates for that setup, as well as in the other setup(s). EncyclopeDIA quantified in total 2,060 *Salmonella* proteins in the 1:1 dilution series, of which 1,091 proteins (53.0%) were quantified in the 1:9 dilution series with a median protein ratio of 0.14 (Figure 4). Furthermore, 264 of these 1,091 proteins (24.2%) were detectable at lower abundance (median protein ratio of 0.016) and even at a ~1,000 fold dilution, 47 proteins were still quantifiable. Note that, as can be expected, few protein quantifications in 1:9999 dilution series were not in line with the expected ratio and were not considered further here. Also, DDA-based MaxQuant quantification reports 16 quantified proteins in the 1:999 dilution series. While showing protein ratios in line with the expected ratios as observed in case of DIA, the number of quantified proteins is considerably lower for DDA analysed samples compared to DIA analysed samples.

**Figure 4.**
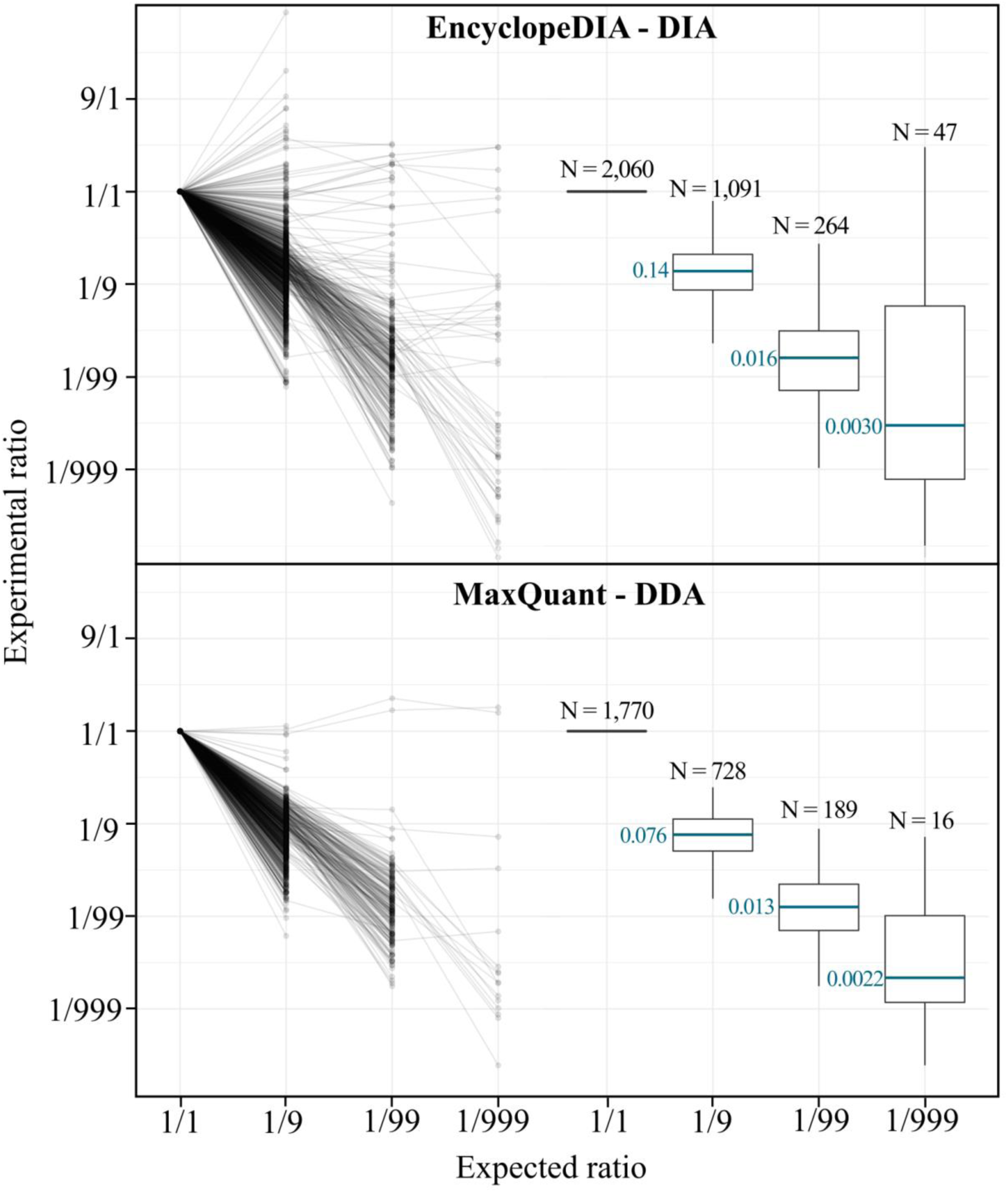
*Salmonella* protein quantification throughout *Salmonella*-human proteome dilutions. Comparison of *Salmonella* protein quantifications obtained using EncyclopeDIA and searching the hybrid library (DIA data, upper panel) and MaxQuant LFQ quantification (DDA data, lower panel). Experimental ratios (y-axis, 9:1 to 1:999) versus expected ratios (x-axis, 1:1 to 1:999) for quantified proteins (left panel) and proteins quantified in at least two out of three replicates among all setups analyzed are indicated with their respective median (right panel).

We also investigated manually the more extreme cases of quantifiable *Salmonella* peptides in the 1:999+ dilutions in our DIA workflow. As a representative example, the peptide ILADIAVFDK (doubly charged) maintains similar MS2 peak shapes throughout 1:1, 1:9 and 1:99 and to a lesser extent in 1:999 and 1:9999 dilutions, while the intensity decreases according the dilution factor (Supplementary Figure 1). However, at increased dilutions, flanking noise peaks are evidenced and become dominant over genuine signalling peaks. As in case of DDA, this represents a challenge for achieving correct identification and quantification in the 1:999 and 1:9999 setups.

## DISCUSSION

MS acquisition of artificial human-*Salmonella* dual proteome mixtures in DIA mode significantly improved identification and quantification of low abundant *Salmonella* proteins compared to DDA acquisition mode, increasing the discriminative power of LC-MS/MS. Among the artificial mixtures (1:1 to 1:9999) used in this study to effectuate comparative analysis between both acquisition modes, the 1:99 dilution closely reflects the host/pathogen ratio in actual infection conditions in case of *Salmonella* [21], where a scarce amount of protein material comes from the bacterial pathogen, making the simultaneous study of host and pathogen proteomes challenging. At a ~100-fold dilution, 98 *Salmonella* proteins were consistently quantified by MaxQuant (DDA) in all three replicates, compared to 279 proteins quantified in all 3 replicates when searching the hybrid DIA/DDA spectral library. Also, in 1:1 *Salmonella*-human proteome mixtures, human and *Salmonella* peptide/protein identification rates were significantly boosted when searching DDA-based spectral libraries. Taken together, the increased identification of peptides with lower abundance in DIA points to the stochastic nature of DDA. Notably, to correct for DDA stochastic sampling, more advanced DDA precursor selection algorithms have been developed such as MaxQuant.Live [42]. In contrast to a recent report [11], *Salmonella* sample pre-fractionation proved to be instrumental to increase *Salmonella* peptide identification, nearly doubling *Salmonella* peptide identification in the 1:1 *Salmonella*-human proteome mixture. This effect is likely attributable to the dual proteome complexity, rendering DDA unable to fully grasp *Salmonella* proteome complexity. Hence, in such multi-species proteome analyses, comprehensive spectral libraries and pre-fractionation of a species of interest can be advisable.

Besides assessing the performance of DDA-based spectral libraries, we tested alternative DIA workflows that enable to search both *Salmonella* and human proteomes. The library-free EncyclopeDIA built-in PECAN algorithm (Walnut) delivered lower peptide identifications, in line with earlier observations [37]. In addition, we made use of MS^2^PIP-predicted spectral libraries, predicting RT with DeepLC trained on DIA identifications made by PECAN. Similar to recent studies reporting on the use of Prosit and Prism-predicted spectral libraries [14, 43], we achieved a similar performance compared to searching DDA-based spectral libraries (Figure 3). Interestingly, both alternative approaches allowed to identify 18,736 novel peptides not found in DDA data. Further inspection of non-DDA peptides identified by using the MS^2^PIP-predicted spectral library, pointed to relatively lower abundant peptides that might have been missed in DDA precursor selection. Notably, the majority of these peptides (85%) matched proteins identified by other peptides in DDA. Combining the strengths of both approaches, we generated a hybrid spectral library containing both experimental DDA data results, and DIA search results (Q-value ≤ 1%) from PECAN and the MS^2^PIP-predicted library. The resulting hybrid spectral library identified ~30,000 peptides in all 3 replicates of concentrated (1:99+) human samples, which is 1.5-fold higher than searching DDA spectral libraries only (~20,000 peptides, Figure 3), and 2-fold higher than a DDA analysis (~15,000 peptides, Figure 2). As such, integrating DIA-based peptide identifications with DDA identifications resulted in drastic increase of peptide identification. One consideration is that the hybrid library is based either on iRT or RT for DDA-based spectra or DIA identification results, respectively. However, given the RT calibration step in EncyclopeDIA, this discrepancy caused no issue. Hybrid spectral libraries were very recently reported to improve proteome coverage [29, 37, 41]. Here, we shown this to greatly facilitate dual-proteome profiling, as such complex proteome mixtures pose an enormous challenge to DDA, making DDA-only libraries far from comprehensive. Taken together, making use of DIA-only approaches, predicted spectral libraries, and perhaps publicly available DDA data or DDA-based spectral libraries, DIA analyses no longer seem to depend on sample-specific complementary DDA runs, although these can further improve the performance, as also shown to be the case in our study. Notably, other DIA-only analyses such as spectral deconvolution algorithms, e.g. DIA-Umpire [20], could further strengthen hybrid libraries.

When judging protein quantification, both MaxQuant (DDA) and EncyclopeDIA (DIA) delivered *Salmonella* protein quantifications in line with dilution series up to 1:999 dilutions. In the infection relevant 1:99 sample, 264 proteins were quantified with a relative median intensity ratio of 0.016 compared to 1:1 equal mixed samples. Notably, MaxQuant also delivers 189 (−30%) quantified proteins, and has overall better accuracy, which is clearer at a 1:999 ratio. Note that we did not include 1:9999 protein quantifications as the few that were found had inconsistent protein ratios. When judging for instance the ILADIAVFDK/2+ precursor that was found across all dilutions (Supplementary Figure 1), it is clear how peptide peaks at a 1:9999 dilution become nearly impossible to distinguish from random noise. However, artificial mixtures do suggest infection-relevant host pathogen conditions with an approximate ~1:100 dilution could be profiled in sufficient proteome depth without prior bacteria enrichment, especially so if using increased LC-MS/MS run time by for instance offline pre-fractionations or the use of parallel DIA runs with narrow *m*/*z* windows in consecutive *m*/*z* ranges as demonstrated in the EncyclopeDIA workflow [37]. Overall, the spectral libraries created and provided as supplemental data online will assist future DIA-based research on *Salmonella* infected epithelial host cells.

## SUPPLEMENTARY INFORMATION

**Supplementary Figure 1.**
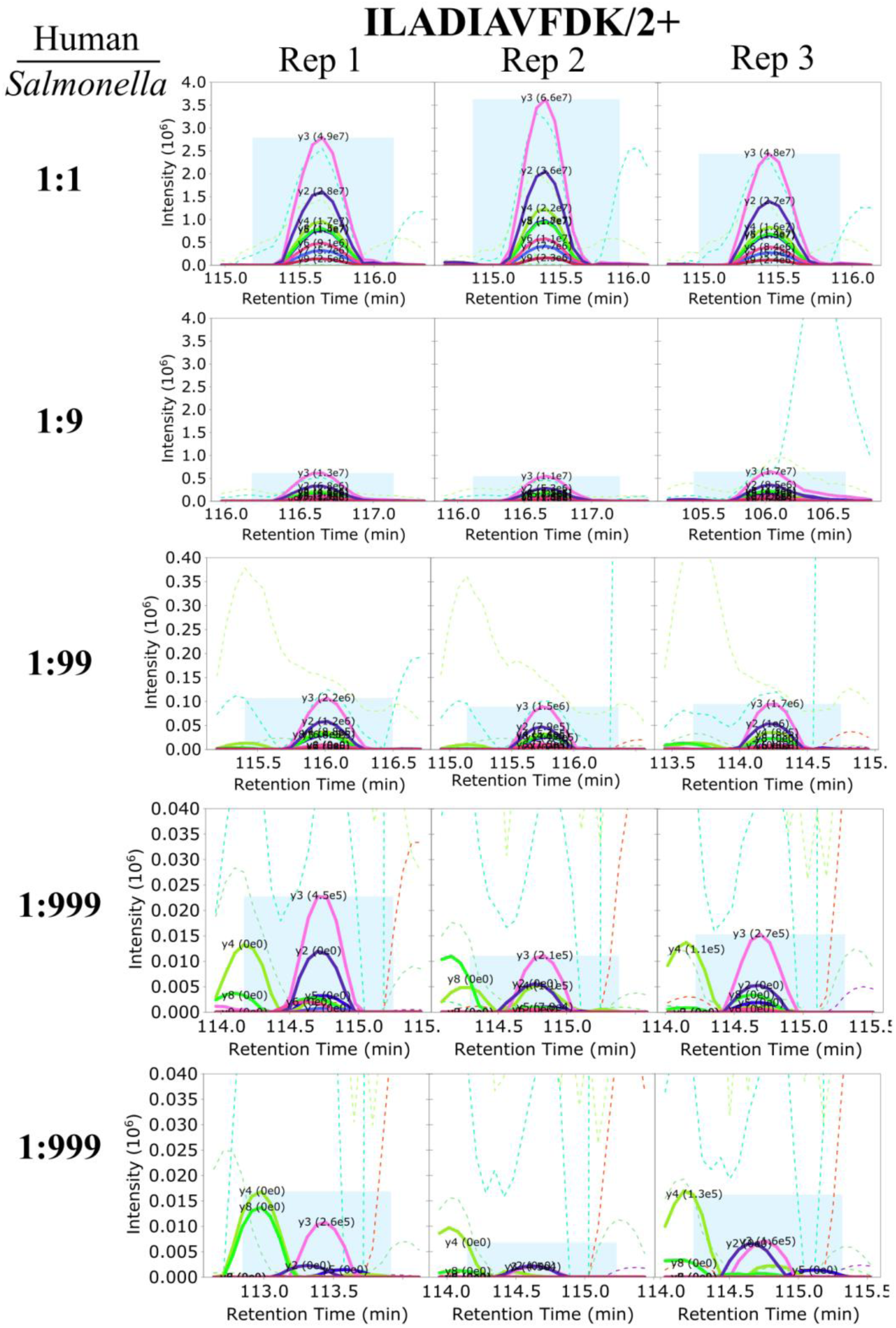
Quantification in DIA workflow of peptide ILADIAVFDK/2+ in replicate samples among the various dilutions (1:1 to 1:9999). Graphics were exported using the multi-ELIB browser and plots MS2 peak intensities (y-axis, different ranges) versus RT (min) (x-axis).

**Supplementary Figure 2.**
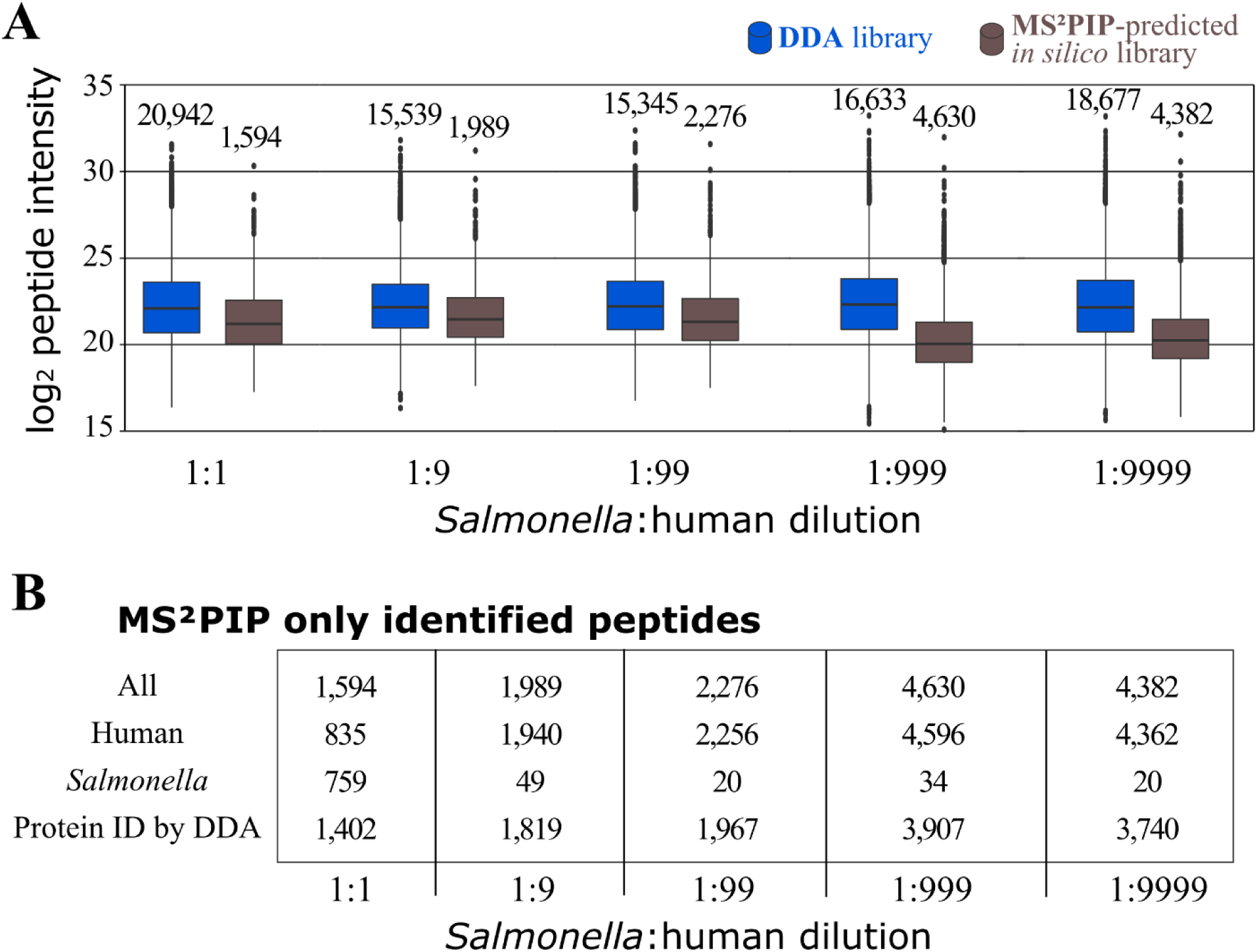
Analysis of unique peptides identified in DIA with the MS^2^PIP-predicted spectral library. **A.** Quantification of identified peptides (y-axis, log2 peptide intensity) across artificial mixtures (x-axis, 1:1 to 1:9999) with the DDA-based spectral library (blue) or with the MS^2^PIP-predicted spectral library. The numbers of quantified peptides corresponding to total identified peptides in the DDA-based spectral library and to novel identified peptides in the MS^2^PIP-predicted spectral library are indicated above each box plot. **B.** Inspection of novel peptides with MS^2^PIP-predicted *in silico* spectral library in relation to human or *Salmonella* origin and by matching to protein entries previously identified in DDA.

### Spectral library and MS data availability

Supplementary data can be found online and the mass spectrometry proteomics data have been deposited to the ProteomeXchange Consortium via the PRIDE [44] partner repository with the dataset identifier PXD017904 (username, password: UeGtkZ2x) for DDA artificial mixtures and *Salmonella* pre-fractionated samples and with the dataset identifier PXD017945 (username, password: 7lArxHj0) for the DIA artificial mixtures. In addition, the generated spectral libraries, FASTA databases, and search results are organized in the Open Science Framework (OSF) project page [45] and are available at https://osf.io/u96bk/.

## AUTHOR CONTRIBUTIONS

P.W., U.F., A.S, K.G. and P.V.D. designed research. P.W., U.F. and P.V.D. performed research. P.W. analyzed data. P.W., U.F. and P.V.D. wrote the paper and all authors contributed to finalizing the manuscript text and gave approval to the final version of the manuscript.

## ACKNOWLEDGMENTS

The authors declare that they have no competing interests. All authors read and approved the final manuscript. We would like to acknowledge Brian Searle for his valuable feedback during the implementation of EncyclopeDIA software.

## ABBREVIATIONS

RT: retention time
iRT: indexed retention time
MS2PIP: MS2 peak intensity prediction

## SYNOPSIS

We performed data-dependent acquisition (DDA) and data-independent acquisition (DIA) on dilution series of artificial dual-proteome mixes of human and *Salmonella* proteomes. Whereas DIA searches of DDA-based spectral libraries led to enhanced recovery of low-abundant peptides in compared to DDA searches, results were further improved by using a hybrid spectral library by including DIA identifications results from a PECAN and an MS^2^PIP-predicted spectral library search.

